# Syntaxin clusters and cholesterol affect the mobility of Syntaxin1a

**DOI:** 10.1101/2023.08.17.552646

**Authors:** Alan W. Weisgerber, Zdeněk Otruba, Michelle K Knowles

## Abstract

Syntaxin1a (Syx1a) is essential for stimulated exocytosis in neuroendocrine cells. The vesicle docking process involves the formation of nanoscale Syx1a domains on the plasma membrane and the Syx1a clusters disintegrate during the fusion process. Syx1a nanodomains are both static yet empty and refill; the process by which these clusters maintain this balance is unclear. In this work, the dynamics of the Syx1a molecules is elucidated relative to the cluster position through a labeling strategy that allows both the bulk position of the Syx clusters to be visualized concurrent with the trajectories of single Syx1a molecules on the surface of PC12 cells. Single Syx1a molecules were tracked in time relative to cluster positions to decipher how Syx1a moves within a cluster and when clusters are not present. Syx1a is mobile on the plasma membrane, more mobile at the center of clusters, and less mobile near the edges of clusters; this depends on the presence of the N-terminal Habc domain and cholesterol, which are essential for proper exocytosis. Simulations of the dynamics observed at clusters support a model where clusters are maintained by a large cage (*r* = 100 nm) within which Syx1a remains highly mobile within the cluster (*r* = 50 nm). The depletion of cholesterol dramatically reduces the mobility of Syx1a within clusters and less so over the rest of the plasma membrane. This suggests that fluidity of Syx1a supramolecular clusters is needed for function.

**STATEMENT OF SIGNIFICANCE:** Syntaxin1a (Syx1a) is essential for exocytosis where the vesicle docking process involves the formation of nanoscale Syx1a domains on the plasma membrane. Syx1a nanodomains are both static yet empty and refill; the process by which these clusters maintain this balance is unclear. In this work, the dynamics of the Syx1a molecules was elucidated relative to the cluster position on the surface of PC12 cells. Single molecules were tracked relative to clusters and Syx1a is mobile on the plasma membrane, more so within a cluster and less mobile near the edges of clusters. This depends on the presence of the N-terminal domain and cholesterol, which are essential for proper exocytosis. This suggests that fluidity of Syx1a clusters is needed for function.

## INTRODUCTION

Many types of membrane proteins form supramolecular clusters to regulate activity^1,2^ or to transmit a signal to the interior of the cell^3,4^. Often the proteins within clusters interchange dynamically with free protein, located outside of a cluster^1,5^, while the clusters themselves appear static^5^. SNARE proteins, the regulators of membrane fusion, form supramolecular clusters to facilitate the fusion of vesicles docked to the plasma membrane^5-7^. The exchange of molecules can affect cluster size and function, thus, the dynamics of membrane proteins and the exchange of free proteins within supramolecular clusters are of great interest as a mechanism of regulating secretion.

In neuroendocrine cells, the secretion of dense core vesicles and synaptic vesicles occurs at sites where Syx1a molecules, a SNARE protein on the plasma membrane, accumulate^6,8^. Several copies of Syx1a are required for fusion^9,10^, yet Syx1a clusters contain 60-80 copies of Syx^1,11^, many more than required. These clusters are stable in their location, yet empty and refill on the order of several seconds^5^. The Syx clusters are thought to form through interactions between the SNARE domain^12^, however, the interactions of the SNARE domain are hindered by the N-terminal Habc domain, which acts as a flap that folds upon the SNARE domain to block interactions with the assistance of Munc18^13^ during Munc18 mediated transport to the plasma membrane^14^. The Habc domain is also required for Syx1a cluster to properly target docked granules^5^. Upon membrane fusion, Syx1a is lost from the fusion site^5,7^, suggesting that the ability of Syx1a to diffuse away during fusion is needed.

The membrane composition also plays an important role in the clustering and dynamics of Syx1a. Syx1a clusters depend on cholesterol^15,16^ and the depletion of cholesterol corresponds to a loss of clusters in PC12 membrane sheets^6^. The loss of cholesterol also reduces vesicle docking and fusion^16-18^. In general, the loss of cholesterol reduces the mobility of plasma membrane protein receptors^19^ and simple transmembrane proteins whose protein-protein interactions are limited^20^, suggesting that cholesterol mediation of protein mobility is a general feature of membranes. Besides cholesterol, Syx1a clusters also recruit PI(4,5)P_2_ by binding through a polybasic region near the transmembrane domain of Syx1a^21^, however this is not required for Syx1a cluster formation nor vesicle docking, but PI(4,5)P_2_ is needed for efficient fusion^22^.

In this work, the mobility of single Syx1a proteins has been measured relative to Syx1a clusters. To measure both cluster and single molecules simultaneously, Syx1a-EGFP was transiently expressed in PC12 cells under a truncated promotor to limit overexpression^5,11^. The cluster positions are evident from Syx-EGFP and single molecules were visualized by sparsely labeling with a red fluorescent nanobody to EGFP^23^. The dynamics of Syx1a near clusters is surprisingly mobile, but this mobility depends on both the presence of cholesterol and the presence of the Habc domain. Loss of cholesterol hindered motion throughout the cell and truncation of Syx1a to remove the Habc domain led to a reduction in motion within clusters. The mobility of Syx1a depends on the cholesterol content and the proximity to a cluster and we propose that the motion of Syx1a within clusters aids in fusion.

## RESULTS

To determine how Syx clusters affect the motion of Syx1a molecules, Syx1a-EGFP was labeled at a low level with a red fluorescent, anti-GFP nanobody (“Syx1a-NB”) and tracked (Schematic in Fig 1A). Syx1a tracks were then compared to the Syx1a-EGFP cluster positions, shown in Fig 1B, and dynamics were determined as a function of the proximity of Syx1a clusters. Syx1a clusters are stable in time over several minutes (Fig 1C) even though they are known to exchange with free Syx1a^5^. Based on this, an imaging protocol was devised (Fig 1D) to measure the single molecule dynamics as they exchange within clusters. Syx1a-EGFP clusters were imaged followed by a 25 second movie of Syx1a-NB (red channel, 500 frames, 50 ms/frame). This was repeated on the same cell, oftentimes, to obtain a movie where the amount of Syx1a-NB was appropriate for single molecule tracking analysis. As expected, the dynamics of Syx1a is highly heterogenous. Example Syx1a-NB trajectories are overlaid on Syx1a-GFP cluster images in Figure 1E-I, where blue in the track marks the start position and red marks the end. Syx1a is mobile and observed as leaving and entering clusters in E-G, however immobile Syx1a molecules are also observed at clusters (Fig 1H) and non-cluster positions (Fig 1I). To measure if the immobile Syx1a molecules were moving slowly or completely immobile, the Syx1a-NB was immobilized on the glass surface, imaged, and tracked for comparison. Fig 1J shows an example track overlaid on the average Syx1a-NB image. Syx1a-NB moves at an average rate of 0.050 µm^2^/s, as measured by the initial slope of the mean square displacement in time of individual trajectories; this is significantly more mobile than the Syx1a-NB stuck to the glass surface (Fig 1K, D = 0.0056 µm^2^/s).

**Figure 1:**
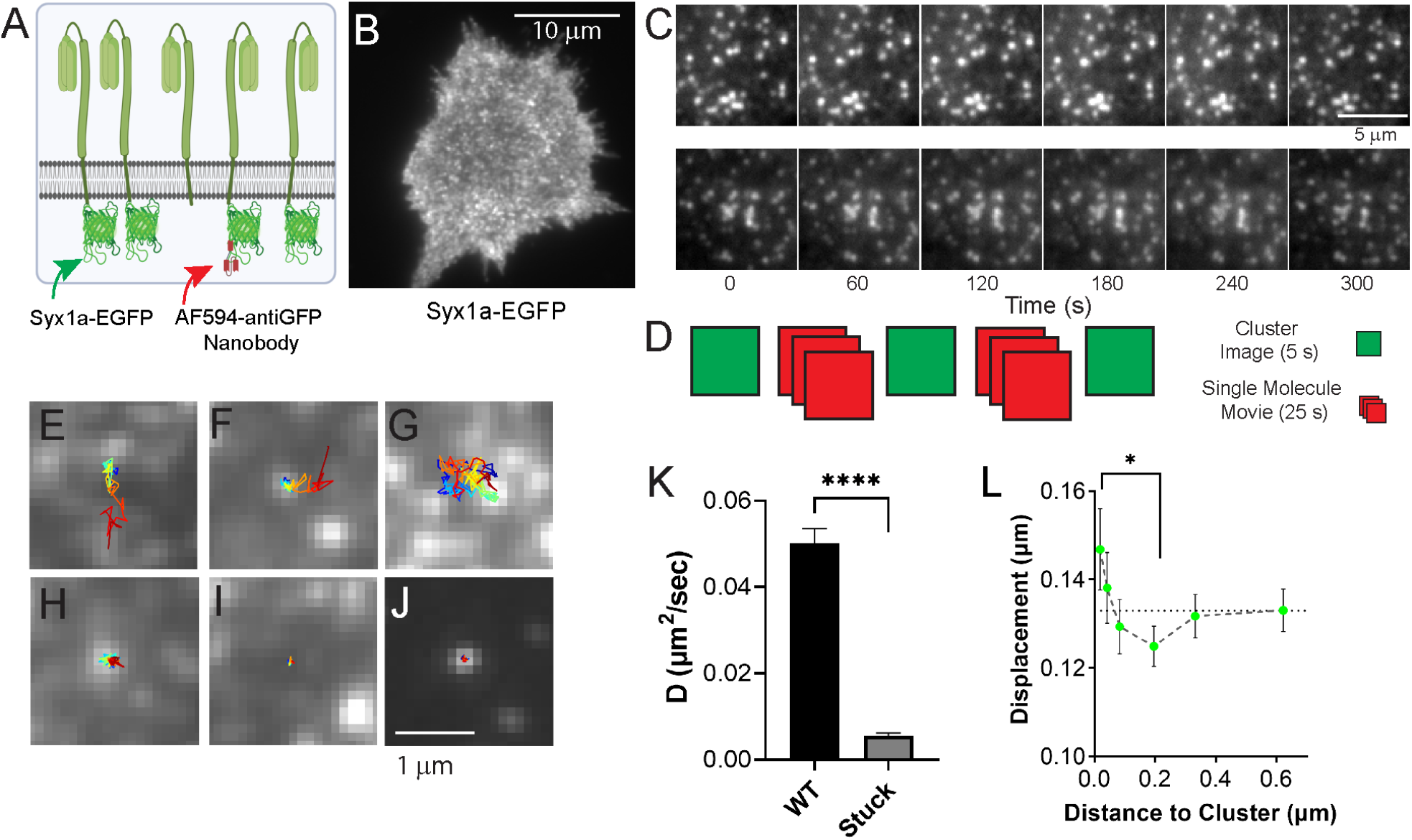
Tracking of Syx1a relative to Syx1a clusters in PC12 cells. A) Schematic for experiment: Cells express Syx1a-EGFP and a dye labeled nanobody (anti-EGFP with Alexa594 dye, “Syx1a-NB”) is added at a low amount to the extracellular space to facilitate single molecule tracking. Syx1a-EGFP is used to locate clusters of Syx1a and the red nanobody is used to track motion relative to cluster positions. B) TIRFM image of Syx1aEGFP forming clusters on the cell surface. C) Clusters were imaged over several minutes to determine if cluster positions are stable in time. Two example cells are shown. D) From this, the imaging protocol was developed where Syx1a-EGFP clusters (green) are imaged followed by a 500 frame (50 ms/frame, red) movie of Syx1a-NB, followed by another cluster image. The cluster images were compared to determine if clusters remain in the same position and the initial cluster image was used to locate clusters. This was repeated 2-4 times. Example Syx1a tracks are shown in E-J and overlaid on the cluster image taken immediately prior to the Syx1a-NB movie. E) Syx1a entering a cluster. F) Syx1a leaving a cluster domain. G) Syx1a entering and then leaving a cluster domain. H) Syx1a stuck at cluster. I) Syx1a stuck not at a cluster. J) Example track of the nanobody, Syx1a-NB, stuck to a glass surface. Tracks start blue and then turn red over the course of time. K) Bulk diffusion coefficient from tracking Syx1a-NB attached to Syx1a-EGFP (N = 21 cells) and the Syx1a-NB stuck to the surface (N = 6 movies). L) The step size over 200 ms of a Syx1a as a function of distance. Syx1a is more mobile at the center of clusters and less mobile 200 nm away. Dashed line indicates the plateau of displacement far from clusters. T-test for significance (p < 0.05).

To assess how single Syx1a molecules interact with clusters of Syx1a, the mobility of Syx1a-NB as a function of distance to the center of the nearest cluster was measured by calculating the step magnitude over a 200 ms interval (4 frames) and the distance between the center of the cluster and the start of the step (Fig 1L). Therefore, one track can contribute multiple steps to Figure 1L. The MSD slope was not used because the distance to the cluster changes throughout the track. The steps were binned as a function of distance from the center of a cluster and the average is shown. Syx1a molecules in the center of the cluster are free to move more than molecules that are a further distance from the cluster center (0.2 µm) and the Syx1a molecules that are furthest from the cluster move at an intermediate rate.

The formation of clusters has been proposed to be due to SNARE domain interactions between Syx1a molecules^12^ and this interaction is limited by the Habc domain^13^. To determine the role of the Habc domain and the SNARE domain, truncated forms of Syx (SyxΔNT and SyxTMD) were imaged (Figure 2). The SyxΔNT construct lacks the N-terminal regulatory domain (NTD) that is known to fold onto the SNARE domain and the presence of the Habc domain reduces SNARE domain interactions^24^. The SyxTMD lacks most of the SNARE domain and the N-terminal domain. If the Syx SNARE domain is responsible for clustering, Syx dynamics should be affected in the form that allows for more SNARE interactions (SyxΔNT) and be reduced in the form that lacks the SNARE domain (SyxTMD). Fig 2A shows the constructs tested. All contain a GFP that resides on the outside of the cell, however, when labeling with Syx1a-NB, no labeling occurred with the SyxTMD (Fig S1). Upon further investigation using confocal microscopy (Fig 2B-D), surface labeling was observed for both SyxWT (Fig 2B) and SyxΔNT (Fig 2C) however the SyxTMD was clearly trapped within the cell (Fig 2D). Both SyxWT and SyxΔNT were able to be labeled by the addition of Syx1a-NB, further supporting that they are indeed on the plasma membrane and SyxTMD is not. Therefore, the tracking studies of SyxWT and SyxΔNT were possible. Clusters were observable in both cases (Fig 2E), but others note that the clusters formed by SyxΔNT do not colocalize as well to docked vesicles as Syx1a clusters^5^, suggesting they are not targeted correctly to potential binding partners. SyxΔNT revealed slightly, but not significantly reduced mobility on the cell surface (Fig 2F). This was consistent with FRAP data that showed a small reduction in the fraction mobile (Fig S2). The mobility of SyxΔNT as a function of distance to a cluster was not significantly different anywhere along the curve (Fig 2G, red), with no hindrance at the border of clusters nor enhancement at the center of clusters as observed with Syx1a-NB dynamics (Fig 2G, green). This suggests that proper targeting of Syx1a, facilitated by the NTD is necessary for the observed cluster based dependent motion.

**Figure 2:**
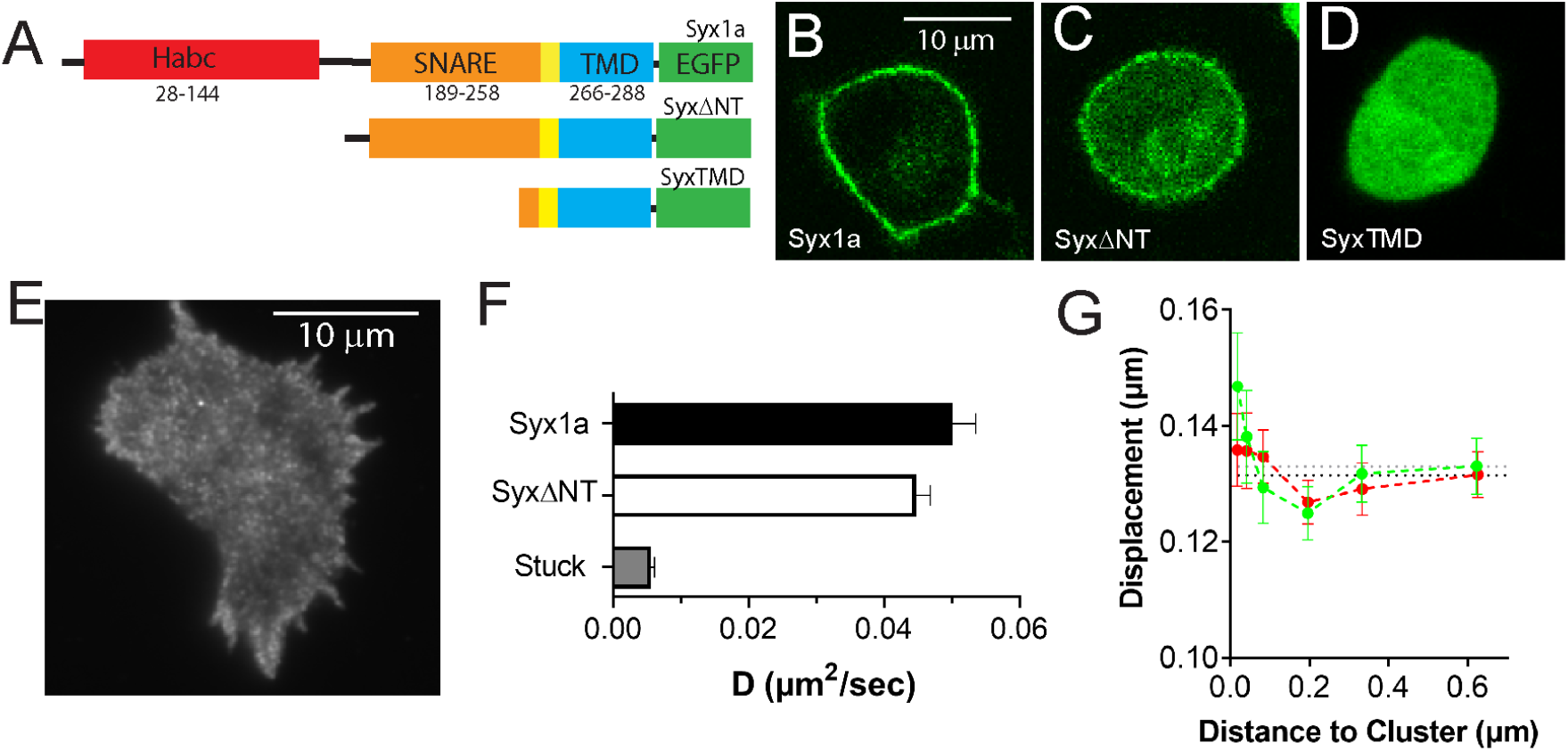
Syx1a lacking the N-terminal domain (Syx1a-dNT-EGFP) was tracked relative to Syx1a-dNT clusters in PC12 cells. A) A representation of the protein domains of Syx1a-GFP, SyxΔNT-GFP, and Syx1a trans membrane domain (SyxTMD-GFP). B – D) Confocal microscopy of Syx constructs imaged halfway through the cell. B) Syx1a-GFP is on the cell surface, C) SyxΔNT-GFP is enriched on the cell surface, but D) SyxTMD-GFP appears throughout the cells. E) Bulk diffusion coefficient from tracked SyxNB, which binds GFP, applied to the extracellular space of cells expressing Syx1a-EGFP (N = 21 cells), SyxΔNT-EGFP (N = 22 cells), and stuck SyxNB (N = 6 movies). F) The step size over 200 ms as a function of distance. Syx1a is more mobile at the center of clusters and less mobile 200 nm away. Dashed line indicates the plateau of displacement far from clusters. T-test for significance shows no significance between the first and fourth distance point (p > 0.05)

To better understand what underlying molecular forces could give rise to the cluster-dependent mobility trends (Figures 1L, 2G), a diffusion-based model was developed and used to simulate the data. The key features in the cluster distance dependent motility observed for wild-type Syx1a (Fig 3A) are the higher mobility at the center of the cluster (Fig 3A, pink zone), the reduced mobility at approximately 0.2 µm from the center of a cluster (Fig 3A, yellow zone), and the mobility when far from a cluster (Fig 3A, teal zone). Here, the cluster size was determined based on published super resolution data (diameter = 93.4 nm)^25^. In this model (Fig 3B), Syx1a molecules move freely at a rate of 0.027 µm^2^/s, to obtain the step-size observed for molecules far from a cluster. To model the reduction in motion around 0.2 µm, a cage exists that has a semi-permeable barrier, where 10% of steps penetrate the border to enter and 2.5% of the steps are able to leave. We propose a cage as opposed to intermolecular interactions because the step size at the center of the cluster is not as hindered. The cage alone can recreate a portion of the observed data (Fig 3C, blue), however, the enhanced motion at the center of the cluster is not replicated unless the mobility of Syx1a within the cluster is allowed to be larger than the mobility in the “far” zone. The enhanced diffusion at the center of the cluster can be simulated by increasing D as a function of distance from the center (Fig 3C, red) and the data can be recreated in a simulation where the rate of motion increases in the center of the cluster and a semi-permeable cage exists (Fig 3D).

**Figure 3:**
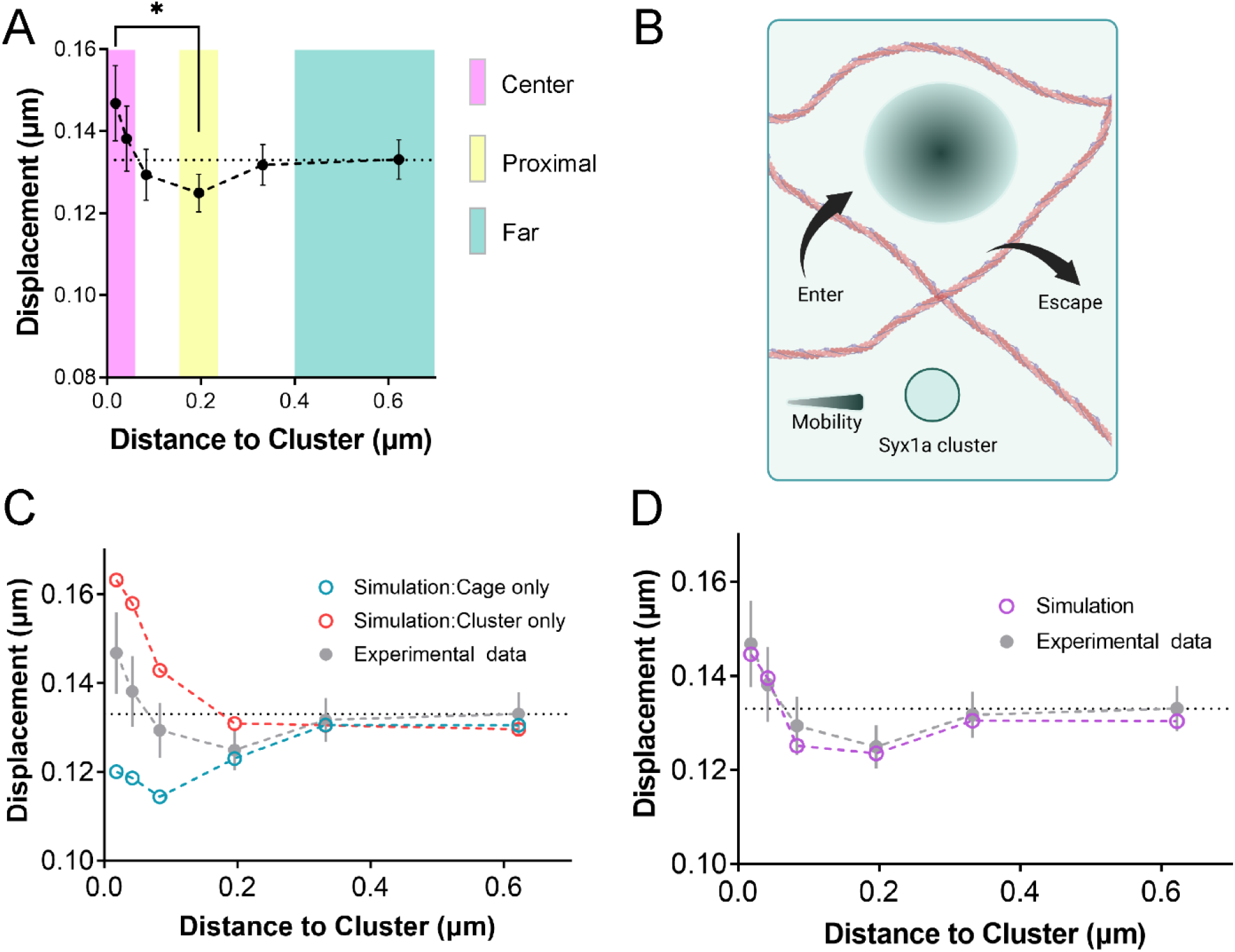
Syx1a dynamics can be modeled with a cage and cluster. A) The displacement of single Syx1a molecules over 200 ms depends on the proximity to a cluster and three zones have been identified. The center of the cluster (pink) exhibits the fastest rate of motion. The proximal zone (200 nm from the center, yellow) exhibits the slowest rate of motion. The molecules far from the cluster (> 400 nm, cyan) move at an intermediate speed and the dotted line notes the mean step size when molecules are far from a cluster. B) To model the dynamics, a cluster (*r =* 50 nm, dark green) was created that has a higher mobility at the center and a cage (*r* = 200 nm) that is semi-permeable (pink). C) Molecules were placed at a certain distance from a cluster and allowed to move 1000 steps with each step taking 200 µs such that the total step is 200 ms. The data is shown in grey and a cage alone (blue) and a cluster alone (red) fail to replicate the data. D) The combination of cluster and cage (purple) can replicate experimental data (grey). All error bars shown are SEM. In simulations, the error bars are smaller than the data points.

Next, we looked to identify molecules that could alter the diffusion coefficient and identified cholesterol as a viable candidate. Cholesterol is well-established in its role to influence the mobility of a wide variety of proteins on the cell surface^19,20,26^, where the depletion of cholesterol reduces the rate of motion by approximately 20-50%^26^. Cholesterol has also been established as important for Syx1a clustering in model membranes^15^, PC12 membrane sheets^6^ and MIN6 cells^16^. Therefore, we hypothesized that Syx1a motion would be reduced when cells were treated with MβCD. To determine if cholesterol is involved with the enhanced diffusion observed within Syx1a clusters, cholesterol was depleted from the membrane. Upon treatment of PC12 cells with 15 mM MβCD, SyxWT moved slower, yet clusters were still visible when compared to untreated cells (Fig 4A-B). Often cells were reduced in size and the rate of diffusion dropped approximately 25% (Fig 4C). This was corroborated with FRAP experiments that show MβCD treatment reduced the diffusion coefficient and the fraction mobile (Fig S2). The dynamics of SyxWT as a function of distance from the Syx1a clusters was measured and compared to cells that were not treated with MβCD (Fig 4D). Interestingly, the motion of molecules near the cluster decreased the most (Fig 4E). To better understand this data (Fig 4D-E), a model similar to the one described above (Fig 3) was used. However, to model the motion of Syx1a when cholesterol is depleted, the cluster dynamics needed to be changed; Syx1a moved slower, not faster, within observed clusters. To reproduce the data, the rate of motion was decreased within the cluster by a factor of 1.2 relative to the surrounding area and was not dependent on distance from the center (Fig 4F). This is analogous to increasing viscosity by a factor of 1.2. All parameters for the model are contained in Table 1. Overall, these data demonstrate that cholesterol is involved with enhancing the mobility of Syx1a within clusters and this likely occurs by altering the viscosity locally. When cholesterol is removed the clusters are less mobile.

**Table 1:** Simulation Parameters. The simulation consisted of freely diffusing molecules that encounter a cage and a cluster. To simulate the cluster, where the molecules increased in their motion when cholesterol was present, the effective viscosity was lowered as a function of distance to cluster, x (µm). Upon cholesterol depletion, the effective viscosity increased within the cluster and was constant. The cage for both simulations was the same and described in the methods.

| Distance from Cluster ($\mu\text{m}$) | Effective viscosity | |
| --- | --- | --- |
| | Syx1a (-MβCD) | Syx1a (+MβCD) |
| 0 to 0.05 | (5x+0.01) | 1.2 |
| > 0.05 | 1.0 | 1.0 |

**Figure 4:**
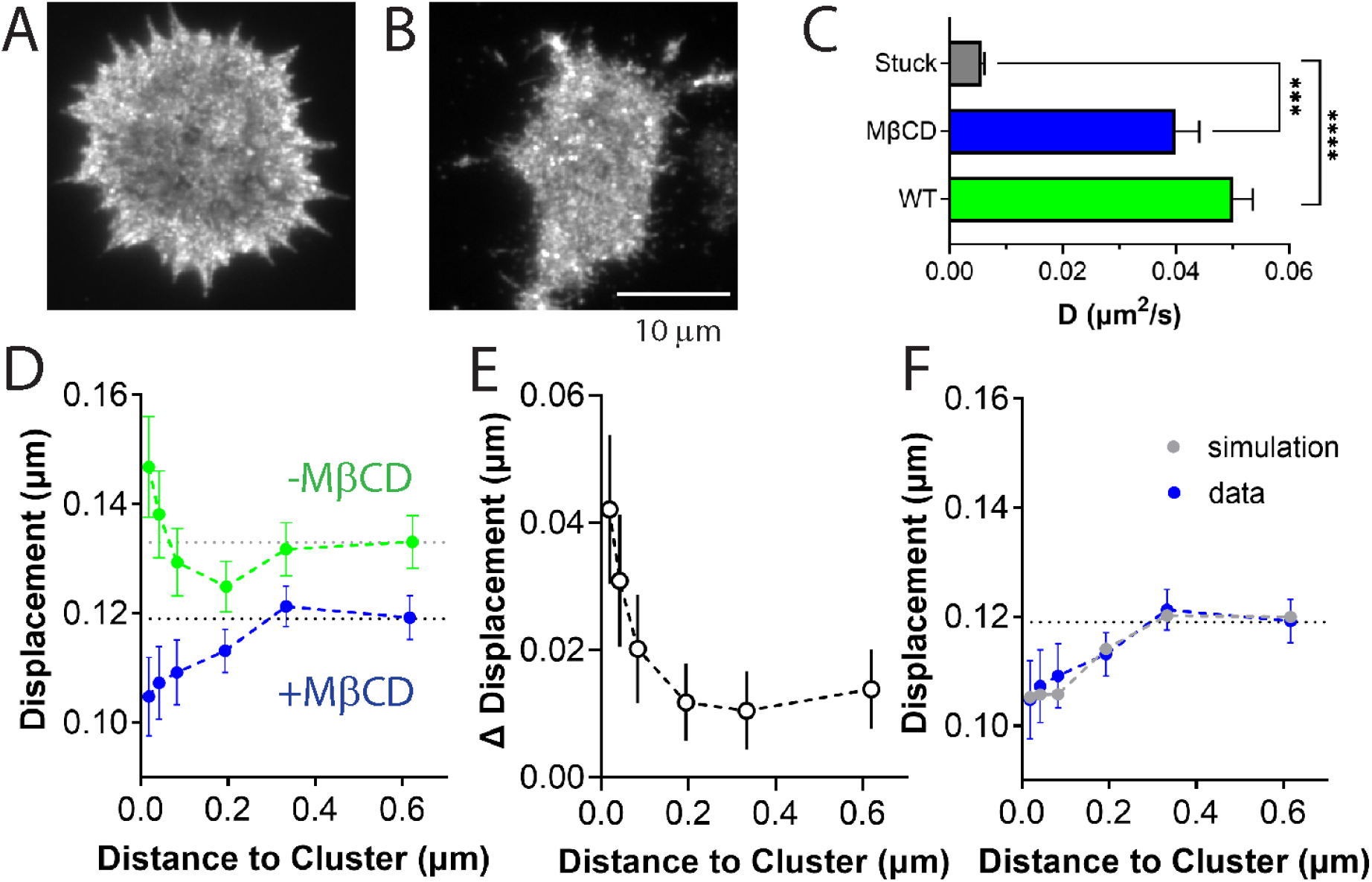
Depletion of cholesterol changes the mobility of Syx1a-EGFP. Cells were treated with 15mM MβCD for 30 minutes prior to imaging. A) TIRFM image of a PC12 cell transfected with Syx1a-EGFP. B) PC12 cell transfected with Syx1a-EGFP and treated with MβCD. C) Bulk diffusion coefficient from tracked AF594NB in Syx1a-EGFP cells without MβCD treatment (N = 21 cells), with MβCD treatment (N = 23 cells) and AF594NB adhered to poly-L-lysine coated wells (N = 6 movies). D) The step size over 200 ms of a Syx1a as a function of distance from a cluster. Dotted horizontal lines indicates the plateau of displacement far from clusters. E) The difference between the steps taken without MβCD treatment (green points in D) and the steps taken with MβCD treatment (blue points in D). F) Simulation of the data with a corral and a constant cluster viscosity.

## DISCUSSION

In this work, we evaluate the dynamics of Syx1a at cluster positions and beyond. Syx1a clusters are well-established as docking and fusion sites for secretory vesicle exocytosis^6,7^. Clusters form beneath secretory vesicles after vesicles approach the plasma membrane and the formation of Syx1a clusters is essential for proper docking^7^. If Syx1a molecules are not recruited to the vesicle location, the vesicle only transiently visits the membrane^7^. Therefore, it is likely that the vesicle contains molecules that interact with Syx1a to establish connections essential for stable docking. Since many copies (50-70) of Syx1a are within clusters^1,11^ it is possible that many molecular connections exist, allowing individual interactions to be weakly associated, but numerous. Past work demonstrated that Syx1a clusters are highly dynamic, with Syx1a leaving and returning to the same position over time^5^, suggesting that the molecules within the cluster are able to come and go as other connections remain. To measure how single Syx1a molecules dynamically interact with clusters of Syx1a, Syx1a clusters were measured concurrently with single Syx1a molecules. This allowed for the dynamics of individual proteins that enter and leave the clusters to be measured. By sparsely labeling externally with a red-fluorescent, anti-EGFP nanobody (NB) and more broadly by transfecting with an EGFP labeled Syx1a for cluster localization, single molecule dynamics at cluster positions can be elucidated (Fig 1A-H). Here, the red fluorescent, anti-GFP NB bound specifically to transfected cells that exposed EGFP on the exterior, which occurs during proper trafficking of Syx1a-EGFP, as shown in our past work^27^. The truncated Syx1a that lacks the regulatory N-terminal domain (SyxΔNT) was trafficked to the surface (Fig 2C), like full length Syx1a (Fig 2B). However, Syx1a that lacked both the N-terminal domain and the SNARE domain was not trafficked properly (Figs 2D and S1). Therefore, only the dynamics of Syx1a and SyxΔNT were measured and compared to cluster positions.

In our data, the motion of Syx1a was dependent upon the cluster positions and the rate of motion was higher near the center of clusters when compared to nearby a cluster (∼200 nm away) or further outside of a cluster (> 0.5 µm away) (Figure 1L, 3A). One limitation of the measurements of clusters here is that the small size of clusters (100 nm) is not resolvable from the TIRF GFP images. Therefore, it is possible that the locations of the clusters identified is a group of smaller clusters^25^. However, the dependence of mobility on a cluster position was unexpected, as a cluster suggests that molecules are relatively stable in their positions and molecular interactions lead to the accumulation of molecules at that location. Therefore, we altered the protein via removal of the N-terminal Habc domain (Figure 2), depleted cholesterol (Figure 4), a known regulator of Syx1a clusters, and developed a model (Figure 3) to further explore the findings.

The N-terminal domain (NTD) of Syx1a has several roles that regulate Syx1a. First, the NTD is required for proper targeting of Syx clusters to docked granules in PC12^5^ and Ins1 cells^7^. This likely takes place through Munc18, where Munc18 binds the Habc domain^13,28,29^, facilitates interactions with docking vesicles^5^ and promotes fusion *in vitro*^30-32^. Second, the NTD binding of Munc18 is also thought to keep Syx in a “closed” state^24^ reducing the exposure of the SNARE domain, possibly to aid in the regulation of SNARE complex formation. Third, the NTD of Syx binds Munc18 to chaperone Syx1a to the membrane^33^. We observe a slight retention of SyxΔNT within the cell in confocal imaging (Figure 2B-C). By removing the NTD, Munc18 is likely no longer bound as strongly to Syx1a^33,34^, the SNARE domain remains more exposed and the targeting to vesicles is hampered. Overall, the mobility of SyxΔNT was less dependent on the distance to a cluster (Figure 2G). There is no significant difference between the mobility of SyxΔNT molecules at the center of clusters when compared to other locations on the cell surface. The removal of the NTD could directly expose the SNARE motif, allowing more SNARE-SNARE interactions with other Syx molecules and this could decrease mobility within clusters.

Although protein-protein interactions play a key role in Syx cluster formation^1,12^, it was difficult for us to envision how SNARE interactions could increase mobility within clusters. Therefore, we looked to the membrane environment, which is known to affect the motion of proteins. Specifically, Syx clusters contain cholesterol^6,15,16,21^ and cholesterol is essential for maintaining fluid cell membranes^19,20^. Although studies on PC12 membrane sheets show clusters are depleted in the presence of MβCD^6^, others have observed clusters remain post cholesterol depletion in whole PC12 cells^35^, similar to our observations (Fig 4A-B). Even in simple, *in vitro*, membranes (POPC liposomes) an increase in cholesterol concentration led to clustering of Syx1a, in contrast to the initially observed dispersed Syx1 molecules^15^. Radiolabeling experiments showed that Syx1a is in close contact with cholesterol^6^ and SNARE proteins are contained in cholesterol-rich lipid rafts in cell fractionation assays^36^. Cholesterol is a key component in cell membranes and makes up 25-50% of the plasma membrane lipids; with Syx1a clusters likely higher in cholesterol concentration due to the direct interactions between Syx1a and cholesterol. Therefore, we hypothesized that cholesterol maintained the enhanced mobility observed at the center of clusters (Figure 1L, 3A, pink zone). By sequestering cholesterol, the mobility of Syx1a was decreased throughout the cell (Figs 4C and S2) and even more reduced within clusters (Fig 4D-E) suggesting that cholesterol is needed for mobility within clusters and elsewhere.

To better understand the observed dynamics at clusters, a model was designed to account for the experimental data (Figures 1L, 3A, 4F) and dynamic simulations were performed. Several models on Syx1a clustering have been previously examined. In one model, Syx1a interacts through the SNARE domains to accumulate into clusters and the NTD folds over the SNARE domain on the periphery of the cluster to restrict entry^1^. This model was designed based on recapitulating FRAP measurements and, to do so, 16% of Syx1a is freely diffusing and the rest is clustered^1^. These experimental results align with our results for Syx1a FRAP (Fig S2). In a second model, *d*STORM was used to characterize Syx1a clusters in PC12 membrane sheets^25^. Dynamics were not measured, but clusters were labeled with antibodies and determined to be 93.4 nm in diameter and asymmetric^25^. Syx1a was denser at the core and closely surrounded by unclustered Syx1a molecules approximately 120 nm from the cluster^25^. A third model used reaction-diffusion kinetics to explain the fast and slow motion of Syx in neurons^37^. In this model, the bound state of Syx fluctuates as it enters synaptic zones and this state alters mobility^37^. In developing our model, we sought to build upon these, taking the cluster size of approximately 100 nm^25^ and the barrier for entry^1^, however, none of the models describe the enhanced dynamics within the cluster observed in the experimental data. Therefore, the model presented here relies on a combination of a cage that restricts entry and exit, and a variable viscosity within the 100 nm cluster. Together, these two features recapitulate the data (Fig 3D) but individually (Fig 3C) they do not. The components of the cage are up for debate, but past work suggests that proteins that bind Syx^14,38^, SNARE complex formation^39^, and the conformation of Syx1a^35^ may all play a role. Future work can elucidate details of these molecular interactions.

The role of cholesterol could simply be to alter the membrane viscosity, however, cholesterol can also affect the cytoskeleton and could play a role in the caging of Syx molecules during cluster formation, as the effects of cholesterol depletion on protein dynamics is challenging to interpret through simple changes in the diffusion coefficient^40^. For example, Rickman *et al* discovered that MβCD treatment disturbed the lipid order and the conformational state of SNAREs, measured by FRET, changed within clusters after cholesterol depletion^35^. Cholesterol can also affect the extent by which a protein sits in the plasma membrane and cholesterol depletion has been shown to reduce the exposure of membrane proteins to the aqueous environment^41^. Consequently, this can affect the mobility. In our work, the rate of motion increases near the center of clusters and the identity of the cage is unknown. However, the cage is needed for simulating experimental data for both Syx1a with (Fig 3F) and without MβCD (Fig 4F), suggesting that the Syx1a dynamic effects due to cholesterol depend on how cholesterol affects the membrane, the extent by which Syx1a sits in the membrane or the conformation of Syx, not the cage itself.

We propose that the mobility of Syx1a within clusters is a key feature needed for proper membrane fusion. Past studies show that Syx1a and Munc18 are rapidly lost from the fusion site, simultaneously with exocytosis^7^. Other studies note the dynamics of Syx1a both during clustering (pre-fusion), where clusters readily and seemingly cooperatively empty^5^, and the rapid loss of Syx clusters post-fusion^5,7^. The mobility of clusters in the motor nerve terminal of *Drosophila*, depended on stimulation, which altered clustering by reducing the size and density of clusters and increasing Syx1a mobility on the cell surface^42^. In this system, the dynamics of Syx was controlled by the disassembly of SNARE complexes by NSF, which increased motion, and the interactions with polyphosphoinositides, which decreased motion^42^. Together, these studies and our results suggest that Syx clusters need to be dynamic and cholesterol is one essential component for maintaining the dynamics of this highly complex supramolecular cluster.

## MATERIALS AND METHODS

### Table of reagents and products used

*Table of reagents and products used:*
| Reagent | Manufacturer (location) |
| --- | --- |
| High glucose DMEM, Fibronectin, Versene | Gibco (Billings, MT, USA) |
| Horse Serum | Hyclone (Logan, UT, USA) |
| Fetal Bovine Serum | Atlanta Biologicals (Flowery Branch, Georgia, USA) |
| KCl | ACROS (Geel, Belgium) |
| MgCl<sub>2</sub>,<br>CaCl<sub>2</sub>,<br>D-Glucose,<br>HEPES,<br>Methyl-β-cyclodextran (MβCD),<br>Na<sub>3</sub>PO<sub>4</sub>,<br>Imidazole,<br>Lysozyme, | Sigma Aldrich (St. Louis, MO, USA) |
| Amicon Ultra-4 Centrifugal Filter,<br>His-Trap HP |  |
| Luria Broth | Alpha Biosciences (Baltimore, MD, USA) |
| IPTG | GoldBio (St. Louis, MO, USA) |
| NaCl,<br>AlexaFluor594 NHS ester,<br>Ampicillin,<br>Alexa594 NHS ester,<br>MycoFluor™ Mycoplasma Detection Kit,<br>Fluospheres (yellow-green, carboxylate modified 200nm),<br>Lipofectamine 3000,<br>Bovine Serum Albumin | ThermoFisher (Waltham, MA, USA) |
| T25 Flasks | Life Science Products (Frederick, CO, USA) |
| 8-well imaging dishes | Cellvis (Mountain View, CA, USA) |

#### Cell Culture

PC12-GR5 cells were cultured in DMEM (high-glucose) supplemented with 5% horse serum and 5% fetal bovine serum and stored at 37°C/5% CO_2_ in T25 cell culture flasks, as described previously^11^. Cell media was changed every two days. For imaging experiments cells were plated on coverslips containing 20µg/mL Fibronectin for a period of 24 hours prior to transfection with Lipofectamine 3000 as described in the manufacturers protocol, using 100 ng of desired plasmid. Cells were then imaged 24 hours post transfection using imaging buffer (140mM NaCl, 3mM KCl, 1mM MgCl_2_, 3mM CaCl_2_, 10mM D-Glucose, 10mM HEPES, pH 7.4). Cells were transfected with the full length Syx1a plasmid (“SyxWT”) or the truncated Syntaxin (180-288)-GFP plasmid that lacked the N-terminal domain (“SyxΔNT”). The SyxWT plasmid, peGFP(N1)-deltaCMV-rSyntaxin1a-meGFP (Addgene plasmid # 34631), and the SyxΔNT plasmid were gifts from Wolfhard Almers^5^. Cells were transfected 24-48 hours prior to imaging. The high expressing CMV promotor was used for FRAP and the lower expressing, delta CMV promotor^5^ was used for TIRF imaging. For MβCD experiments, cells were subjected to 30 min of incubation with DMEM containing 15mM MβCD. Cells were washed 3x with imaging buffer prior to imaging. PC12 cells tested negatively for mycoplasma using MycoFluor™ Mycoplasma Detection Kit.

### Nanobody Purification and Labeling

Anti-EGFP nanobody plasmid (pOPINE GFP) was a gift from Brett Collins (Addgene plasmid #49172)^43^ and transformed into BL21 cells for purification. Starter cultures (10 mL) were made in LB media with ampicillin (100 µg/ml) and used to inoculate 1L culture the next day. Cultures were grown at 37°C until the OD600 reached 0.9, then temperature was changed to 20°C following induction with IPTG (1mM). The culture was given 24 hours to express the anti-EGFP nanobody, after which the bacteria was pelleted with a centrifuge at 6000 x *g* for 15 min at 4°C. Pellets were stored at −80°C until purified. For purification, pellets were thawed on ice and resuspended in 20 mL lysis buffer (300 mM NaCl, 50 mM Na_3_PO_4_, 5 mM Imidazole, pH 8.0) and Lysozyme (1 mg/ml) was added. The lysate mixture was sonicated for 2 minutes and centrifuged at 14,000 rpm for 15 min at 4°C. The resulting supernatant was then filtered using a 0.22 µm syringe filter prior to being purified on the FPLC with a HisTrap HP column. The resulting fraction of interest was dialyzed with 0.1M Sodium Bicarbonate Buffer (pH 8.3) for concentration quantification and dye labeling.

Dye labeling with AlexaFluor594 was accomplished with AlexaFluor594 NHS ester from Thermo Fisher Scientific (A20004) and was done following the manufacturer’s protocol, as described previously^27^. The nanobody was concentrated to 1 mg/mL for labeling using Amicon Ultra-4 Centrifugal Filter. After labeling was completed, nanobody was separated from free dye using Amicon Ultra-4 Centrifugal Filter. Labeled nanobody was stored at 4°C until use.

### Total Internal Reflection Fluorescence Microscopy and Confocal Microscopy

For TIRF microscopy experiments, a Nikon TI-U inverted microscope was equipped with a 491 nm/561 nm laser launch (Solamere Technology Group, Utah, USA) and a DualView (Optical Insights, Exton, PA, USA) that splits the red and green fluorescence (565LP dichroic with 525/50 and 605/75 emission filters, Chroma Technologies) into separate channels onto an EMCCD (Andor iXon897), as previously described^44^. Micromanager software was used to run the microscope^45^. For single molecule tracking experiments, images were first acquired for the green channel, followed by a movie (50 ms/frame, 1000 frames) taken in the red channel. The green channel was used to locate Syx1a-GFP cluster positions; the red channel was used for tracking Alexafluor594-labeled, anti-GFP nanobody attached to Syx1a-EGFP. The red and green images were aligned using 200 nm carboxylate modified, yellow-green fluospheres and a home-built MATLAB code. For imaging experiments cells were plated on coverslips containing 20μg/mL Fibronectin for a period of 24 hours prior to transfection. After 24-48 hours of transfection, samples were blocked with 10% BSA enriched DMEM and allowed to incubate with this solution for 15 minutes, after which samples were incubated with 0.013 µg/mL AlexaFluor594 anti-EGFP nanobody for 15 minutes. Samples were then rinsed 3 x with imaging buffer prior to imaging. For MβCD experiments, 15 mM MβCD was added to all steps. To measure if Syx constructs were trafficked to the plasma membrane, confocal imaging was performed on a point-scanning confocal microscope (Olympus Fluoview 3000). GFP labeled Syntaxin constructs were illuminated with 488 nm excitation and the focus was adjusted to be at a midpoint of the cell in the z-direction. Single images were taken to qualitatively assess the membrane localization.

### Image Analysis

Syx cluster positions were located on a bandpass filtered image and tracking of nanobody was performed as described previously^46,47^. The maximum distance a nanobody could move over one frame was 7 pixels, where one pixel is 0.107 µm. Only tracks that were longer than 15 frames (50 ms/frame) were used for subsequent analyses. Cluster positions were identified within single images taken prior to the tracking movie (25 s long) with the same spot finding algorithm. We used freely available code from a Matlab particle tracking repository^48^ and Matlab version R2020a (MathWorks, Natick, Massachusetts, USA). The diffusion coefficient was calculated from the individual mean square displacement (MSD). The initial four time points (50 ms/frame) were fit with a line and the slope divided by 4 is the reported diffusion coefficient.

The resulting tracking data along with cluster localizations were used to calculate the step size relation to cluster locations. Mobility was measured as a function of distance to the nearest cluster by first calculating step sizes (dt = 200 ms, 4 frames) and pairing that with the distance to the nearest cluster from the beginning of the step. The steps for all tracks were compiled and binned according to distances (bin edges were: 0, 0.25, 0.5, 1, 2.5, 3.7, 256 pixels). This was done for each cell. The cells were then averaged and the SEM was calculated from cell to cell. For all Syx1a tracking data, 21 cells from 3 days were analyzed. For SyxΔNT 22 cells from 3 days were analyzed and for Syx1a treated with MβCD 23 cells from 3 days were analyzed. Syx1a control cells were measured on the same days as ΔNT and with the MβCD treated cells.

To calculate the precision in our single molecule tracking measurements, Alexa594-NB was adhered to a poly-L-lysine coated surface and imaged as described above. To determine the localization precision, the standard deviation of the position was calculated and precision was measured according to past work in the super-resolution field^49^. The precision in x and y was the same and was 39 nm (n = 3 movies and 2169 tracks).

### Modeling of cluster dynamics

To simulate the experimental data, 10,000 Syx molecules were deposited at a specific and known distance from a cluster, ranging from 0 to 700 nm from the cluster position. The molecules were allowed to diffuse at a rate of 0.027 µm^2^/s (Syx1a) or 0.023 µm^2^/s (Syx1a + MβCD) taking 200 µs steps for 1000 steps, corresponding to 200 ms. The diffusion rate was determined by running the simulation for molecules placed 0.62 µm from the cluster, where no cluster interactions typically occur, and then matching the step size observed in experimental data at long distance (Fig 3A, dotted line). After molecules travel for 200 ms, the final position was measured, and the total distance traveled calculated. Localization error was included in the final position by adding or subtracting a random value from a normal distribution centered around 0 with the standard deviation of 0.05 from both the x and y position. The mean of the magnitude of this distribution is 39 nm, the experimental precision of the tracking measurements. The data obtained from the simulation is the mean distance traveled over 200 ms for the 10,000 molecules and the starting distance from cluster. The simulation was run 5-7 times for each data set (Fig 3C-D, Fig 4F) and error bars (SEM) were calculated for the variability between simulations; these are typically smaller than the data points in all plots. During the simulation, when molecules were approaching a cluster, two features that affect dynamics were encountered (shown in Fig 3B): 1) a cluster (radius = 50 nm) located within 2) a larger, semi-permeable cage (radius = 0.2 µm). The cluster size was taken from super-resolution data of Syx clusters within fixed cells^25^. The cage radius was determined from experimental data, as the distance where Syx1a exhibited the lowest mobility (Fig 3A, yellow). To recapitulate the experimental data for Syx1a, the cluster had a variable viscosity where the cluster was less viscous at the center (see Table 1 for the function that was used). The specific function did not drastically affect results and we chose the simplest - a linear function. The sizes of clusters and cages were fixed in all simulations. When a particle meets the cage from within the cluster, it can either pass through or be deflected based on an escape probability, a simulation parameter that is determined by a best fit of the data. Analogously, when a particle hits the corral from the outside, its fate is governed by an entry probability, a simulation parameter. The escape and entry probabilities were determined as 2.5% and 10% respectively. We note that the ratio of these probabilities had a much stronger impact on the results of the simulation than their absolute values. To fit the experimental data, the effective viscosity was determined by the first three points in the plot because these are the ones within the cluster, the cage entry/exit probability ratio was determined by the midpoints. The D was determined by the 6^th^ point in the data set, the point furthest from the cluster. This was not fit but calculated by converting dx to D mathematically.

All data was analyzed for statistical significance and figure making in GraphPad Prism.

## Acknowledgements

This work was funded by the National Science Foundation Division of Molecular and Cellular Biosciences (Grant # 2122289).

## Notes

### Competing Interest Statement

The authors have declared no competing interest.

