## Supplementary Figures for "Syntaxin clusters and cholesterol affect the mobility of Syntaxin1a"

### Supplemental Methods:

*Fluorescence Recovery After Photobleaching (FRAP):*

PC12 cells were transfected for 24-48 hours with Syx1a-EGFP or SyxdNT-EGFP then transferred to imaging buffer, as described in the main paper. An Olympus Fluoview 3000 was used to image and bleach the GFP labeled Syx proteins. A 3.98 um circular region was bleached using 488 nm excitation. Three images were collected prior to bleaching to measure photobleaching inherent to the imaging process and data was corrected for this bleaching as described in our past work. Images were collected after intentional photobleaching and fit as described previously[1] to determine the diffusion coefficient and fraction mobile. FRAP was measured at 20-22°C. Graphpad Prism was used for all fitting, t-testing and plotting of data.

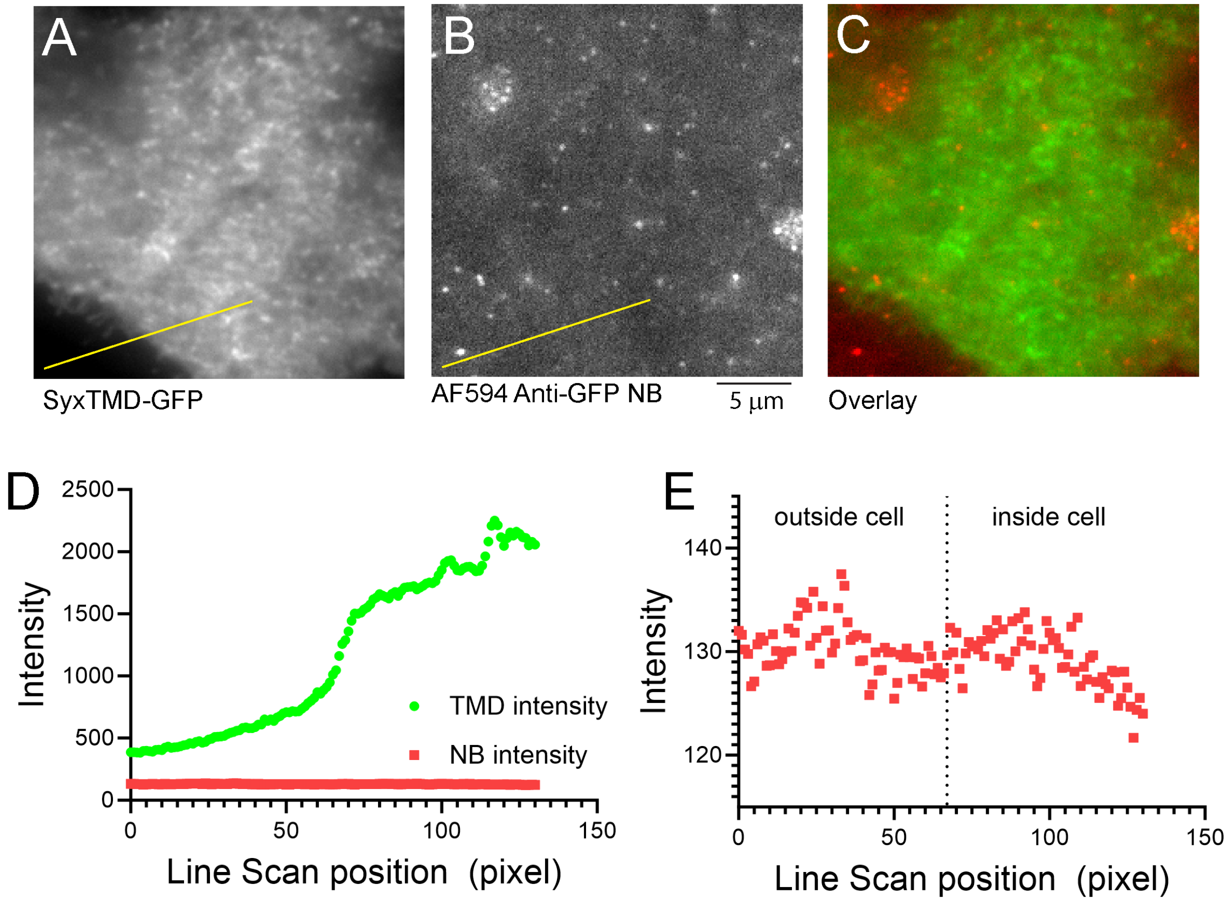


**Supplemental Figure 1: The AF594 labeled, anti-GFP nanobody does not bind cells transfected with SyxTMD.** A) PC12 cells were transfected with SyxTMD-GFP and imaged using TIRF microscopy. An average image is shown. B) Cells were incubated with the AF594 anti-GFP NB that binds GFP if GFP is on the outside of the cell. An average TIRF image (n = xx frames) is shown and the intensity autoscaled. C) An overlay of A and B. D) A line scan of the lines drawn in A and B. The TMD intensity increases at the cell border but the AF594 Anti-GFP NB does not. E) A zoomed in view of the AF594 Anti-GFP NB intensity along the line scan.

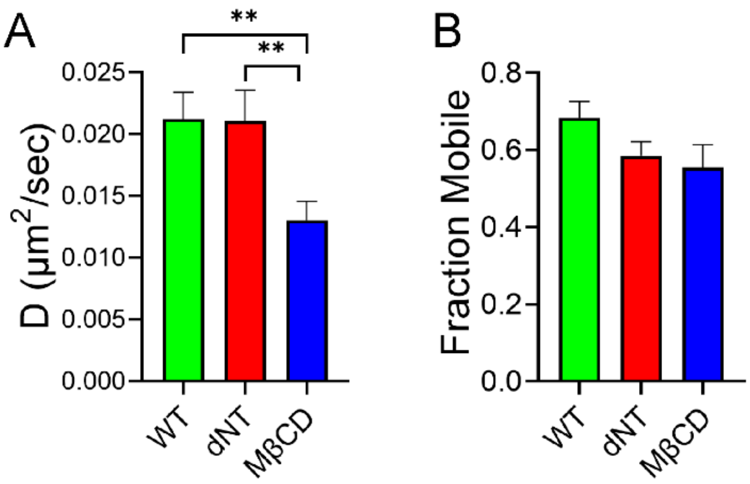


**Supplemental Figure 2: FRAP of PC12 cells expressing Syx1a or SyxΔNT with and without addition of MβCD**. PC12 cells were transfected with Syx1a-EGFP or SyxΔNT and then a 1.99 µm radius circle was photobleached. The recovery was fit according to Eq. 1 to obtain A) Diffusion coefficients and the B) fraction mobile. Syx1a (n = 16 cells), ΔNT (n = 12 cells), and Syx1a cells treated with MβCD (n = 14 cells) were measured. Cells treated with MβCD were significantly slower than untreated Syx1a cells (unpaired t-test, p-value = 0.0060) and SyxΔNT cells (unpaired t-test, p value 0.0088). There is no significant difference between Syx1a and SyxΔNT. The fraction mobile (B) did not test significantly different.
